# Nitro-fatty acids formation in Virgin Olive Oil during gastric digestion and its relationship to cultivar and fruit ripening

**DOI:** 10.1101/592147

**Authors:** Beatriz Sánchez-Calvo, Mauricio Mastrogiovanni, Paula Conde-Innamorato, Mercedes Arias-Sibillotte, Andrés Trostchansky, Homero Rubbo

## Abstract

Virgin olive oil (VOO) represents the main source of unsaturated lipids in the Mediterranean diet associated with low mortality. Health benefits of VOO rely on its composition, mainly fatty acids and minor components such as polyphenols. In addition, VOO contains nitro-fatty acids (NO_2_-FA), novel signaling mediators exhibiting pleiotropic anti-inflammatory responses. Previous work from our group reported the presence of nitro-oleic acid (NO_2_-OA), nitro-linoleic acid (NO_2_-LA) and nitro-conjugated linoleic acid (NO_2_-cLA) in extra virgin olive oil under gastric conditions. Herein, we analyzed the fatty acid profile, phenol, pigment and NO_2_-FA formation in two contrasting Uruguayan olive cultivars, Arbequina and Coratina at two ripening conditions. We demonstrate that VOO fatty acid nitration is dependent on olive cultivar as well as fruit ripening. Under gastric nitration conditions, the presence of polyphenols in Arbequina VOO promoted fatty acid nitration. In contrast, the absence of polyphenols favor lipid oxidation, decreasing fatty acid nitration. In Coratina, where the content of polyphenolic compounds is higher than in Arbequina, their absence did not affect the formation of NO_2_-FA. Coratina contains other bioactive constituents such as pigments that could play an important role in protection of VOO from lipid oxidation. Overall, we postulate that unsaponifiable constituents of VOO, e.g. polyphenols and pigments, contribute to the formation of NO_2_-FA in gastric conditions, thus potentiating their health beneficial

## Introduction

Virgin olive oil (VOO) has increased its consumption worldwide as a lipid component in diets of high nutraceutical value. According to IOC data [1], the total world consumption of olive oil increased by 55% between 1990 and 2000 and an additional 25% between 2000 and 2010, remaining constant until 2018. This represents an incremental difference of 80% in the last 28 years. This increase responds to the knowledge generated about the health benefits of olive oils, which are attributed to its composition and the physical extraction process distinguished as a natural fruit juice [2,3,4,5,6,7,8,9,10,11]. The saponifiable fraction represents 98% of the oil weight being oleic acid (18:1, OA) the fatty acid in greater proportion between 55 and 83%. Linoleic acid (18:2, LA) can reach 20% and linolenic acid (18:3, LnA) up to 1%. The main saturated fatty acid is palmitic acid (16:0, PA) with percentages between 7.5 and 20%, followed by stearic acid (18:0, SA) from 0.5 to 5% [1]. The unsaponifiable fraction, which represents approximately 2% of the total weight, includes chlorophylls, volatile compounds, carotenoids and phenols, the latter being the main antioxidant components in VOO. These compounds also define sensory quality, bitterness, spiciness, aromas and are associated with human health benefits [3,7,8,12,13,14,15,16]. Moreover, VOO contains a high monounsaturated fatty acids (MUFA)/ polyunsaturated fatty acids (PUFA) ratio that confers oxidative stability, improving its conservation.

Multiple health benefits are linked with diets rich in olive oil, including a decrease in cholesterol levels and low density lipoprotein oxidation as well as a decrease in inflammation, endothelial dysfunction, blood pressure, thrombosis and hyperglycemia, among others [17]. Notably, the Mediterranean diet is also linked with the consumption of vegetables that are rich in nitrite (NO_2_^-^) and nitrate (NO_3_^-^) [18,19]. Recent data from our group showed the presence of endogenous nitro-fatty acids (NO_2_-FA) in fresh olives and extra virgin olive oil (EVOO) as well as their formation when EVOO is exposed to artificial gastric fluid [20]. In fact, the acidic milieu of the gastric compartment in the presence of NO_2_^-^ promotes a nitrative environment that can mediate fatty acid nitration [21,22]. Nitro-fatty acids are formed through reactions of nitric oxide-derived oxidant species, e.g. nitrogen dioxide (•NO_2_), with either free or esterified PUFAs [23,24,25]. While under basal conditions the vast majority of the NO_2_-FA detected corresponds to nitro-conjugated linoleic acid (NO_2_-cLA), under gastric conditions the formation of nitro-oleic acid (NO_2_-OA), nitro-linoleic acid (NO_2_-LA) and NO_2_-cLA significantly increased, being 9-NO_2_-cLA and 12-NO_2_-cLA the most prevalent [20].

The formation of NO_2_-FA is linked to cytoprotective and anti-inflammatory signaling responses [26], including inhibition of leukocyte and platelet activation [27], vascular smooth muscle proliferation [28],lipopolysaccharide-stimulated macrophage cytokine secretion [29] activation of peroxisome proliferator-activated receptor-γ [30,31] as well as induction of endothelial heme oxygenase 1 expression [32]. In particular, NO_2_-OA has been shown to play protective roles in hypoxia-induced pulmonary hypertension [33], diabetic nephropathy [34], amyotrophic lateral sclerosis [35] and Ang II-induced hypertension [36] disease models.

It is well known that VOO fatty acid composition, phenolic content and organoleptic properties strongly depends on olive cultivar, fruit ripening, fruit load, hydric availability, edaphic and climatic conditions, productive systems [37,38,39,40] and oil extraction method [41]. Uruguay is situated between 31 and 34 latitudes, analogue of Mediterranean region, with different edaphic and climatic conditions, which implies productivity challenges. Despite this, Uruguayan olive oils have obtained important international awards. Oil composition of Arbequina and Coratina, two main cultivars in Uruguay, are contrasting, having Arbequina olive oil lower polyphenol and OA content than Coratina olive oil [42,43,44,45]. Olive oil quality is defined by international regulations of both the European Community, the International Olive Council [1] and the Codex Alimentarius, which define a series of physicochemical and organoleptic parameters for the identification of different categories. These aspects guarantee oil genuineness, its extraction process and oil conservation. The profile of fatty acids, the contents of polyphenols and pigment represent key tools to describe the nutraceutical value of olive oil. Developing a new nutraceutical factor, i.e. the formation of NO_2_-FA linked with the consumption of vegetables, would be a way to add value to the product, especially, taking into account the well-documented protective actions of NO_2_-FA in human health. In order to achieve this, we analyzed a) the fatty acid profile, phenols, pigments and NO_2_-FA formed in two contrasting Uruguayan olive cultivars, Arbequina and Coratina, at two ripening conditions; and b) the cultivar and fruit ripening effects on NO_2_-FA formation under gastric conditions.

## Materials and methods

### Materials

All chemicals were purchased from Sigma-Aldrich (St. Louis, MO, USA) if not stated otherwise. Nitro-fatty acids internal standards NO_2_^-^[^13^C]OA and NO_2_^-^ [^13^C_18_]LA were obtained from Bruce Freemans’ lab from University of Pittsburgh, USA. Strata NH_2_ (55 µm, 70A) columns were from Phenomenex (8B-S009-EAK).

### Olive oil

Fruits from two olive cultivars, Arbequina and Coratina, growing at an intensive rainfed commercial plantations on east Uruguay (34° 37’ S 54° 36’ O), were harvested at two different growing times, early and intermediate stage (ES and IS), as measured by their maturity indexes (around 1 and 2.5, respectively) [46]. Cold-pressed olive oil was obtained with an Oliomio two phase equipment installed at INIA Las Brujas Research Station–Uruguay. After filtering, oil samples were kept in brown glass bottles at 4 °C for total phenolic content and fatty acid profile analysis; or frozen at −80 °C for NO_2_-FA analysis.

### Determination of phenol content

Total phenol content was determined using Folin-Ciocalteau method described by McDonalds *et al.* [14], with minor modifications. Olive oil samples (5 g) were dissolved in 7 mL of MeOH:H_2_O (80:20, v/v) and vortexed. The mixture was centrifuged for 10 min at 5800 rpm and the supernatant collected and extracted again twice. Afterwards, the collected solution was brought up to volume of 25 mL with MeOH:H_2_O (80:20, v/v). An aliquot (1 mL) was transferred to a 10-mL volumetric flask to which 8 mL distilled water was added. Folin-Ciocalteau reagent (0.5 mL) and saturated Na_2_CO_3_ (0.5 mL) were then added simultaneously and the sample shaken and left for 15 min at room temperature in dark. The absorbance was determined spectrophotometrically at 760 nm in a UV–VIS spectrophotometer (Shimadzu™ UV-3000).The amount of phenols was calculated and expressed as mg of equivalent of gallic acid · kg^-1^ of oil by using a calibration curve prepared with gallic acid.

### Determination of pigments

Chlorophylls (CHLs) and carotenoids (CARs) content were determinated from absorption maximum from an oil solution in cyclohexane at 670 and 470 nm, respectively. The absorptivity coefficients of pheophytin “a” and lutein were used for CHLs and CARs respectively, and the results were expressed as mg/kg of olive oil [15].

### Fatty acids profile

Fatty acid methyl esters were prepared from a trans-esterification reaction with a cold methanolic solution of potassium hydroxide and analyzed by gas chromatography (GC) (Standard IOC/T.20/Doc. n° 24 2001) on a Shimadzu™ GC equipment (Model 14 B) according to [47].

Fatty acids profile was analyzed under gastric conditions in two olive cultivars and two maturation stages: early (ES) and intermediate (IS) stage. Olive oil (10 µl) was incubated for 1 h at 37 °C in 1 mL of gastric juice (HCl 79 mM and NaCl 34 mM) with 0.5 mM NaNO_2_ under continuous stirring [20]. The lipid fraction was extracted with hexane as previously [14], vacuum dried and 1 mL pancreatic lipase (5 mg protein/mL) and phospholipase A_2_ (PLA_2_, 40U) in 50 mM phosphate buffer, pH 7.4 were added, and the reaction mixture incubated at 37 °C for 1 h under continuous agitation. Alternatively, lipids were extracted by Bligh and Dyer [48], vacuum dried and dissolved in chloroform. Lipid classes were further separated by solid phase extraction (SPE) using Strata NH2 columns (100 mg /1 mL). Briefly, columns were pre-conditioned with 1 mL of MeOH followed by 1 mL of chloroform; samples were added and the column washed with 1 mL of chloroform. Free fatty acids were eluted with 1 mL diethylether/2 % acetic acid. The solvent was evaporated under vacuum and lipids were dissolved in MeOH for HPLC-ESI-MS/MS and high resolution mass spectrometry analysis.

### Detection and characterization of NO_2_-FA

Samples were analyzed using a hybrid triple-quadrupole linear ion trap mass spectrometer (QTRAP 4500, AbSciex) coupled online with a liquid chromatography system (Infinity 1260, Agilent). Standard solutions of NO_2_-FA were separated using a reverse phase HPLC column (5µm, 150 × 2.0 mm; Phenomenex) and optimization of mass spectrometer parameters was performed. The electrospray voltage and declustering potential were set to −4.5 kV and −30 V, respectively and the source temperature was 400 °C. Multiple Reaction Monitoring (MRM) analysis was performed for detection of NO_2_-FA; transitions used were *m/z* 326/46 and *m/z* 326/279 for NO_2_-OA, *m/z* 324/46 and *m/z* 324/277 for NO_2_-LA and NO_2_-cLA, *m/z* 344/46 for NO_2_-[^13^C_18_]OA and *m/z* 325/47 for [^15^N]O_2_-cLA. Data were acquired using Analyst 1.6.2 software (ABSciex) and analyzed using MultiQuant 2.1 (ABSciex).

### Statistical analysis

The results are expressed as mean ± standard deviation of the means with values from at least three independent experiments and analyzed with GraphPad Prism®(version 5.0). Statistical analyzes were performed by analysis of variance (ANOVA) followed by Tukey post-test. Differences were significant when p ≤ 0.05.

## Results and Discussion

### Effect of cultivar and rip stage in fatty acid composition

In this study, greater levels of OA as well as MUFA/ PUFA ratio were recorded in Coratina olive oil compared to Arbequina. The content of OA in Arbequina olive oil decreased almost 3.4 % at IS, while LA content increased 1.5 % (Table 1). In Coratina olive oil, OA remained constant at both stages but LA decreased 1.5 % on IS (Table 1). Palmitic acid was higher in Arbequina but in both cultivars decreased within the rip stage. The MUFA/PUFA ratio in Coratina increased at IS while decreased in Arbequina (Table 1). This is in agreement with the results reported on eighteen varieties of Mediterranean origin, including the two varietals analyzed in this study, where three different groups of oils were proposed based on MUFA/PUFA ratio [49]. Herein, Coratina oil was included in group 1 showing high MUFA/PUFA ratio (6.29–21.53 %), while Arbequina oil was included in group 3 having low MUFA/PUFA ratio (1.72–5.50 %), being richer in PA (16.32–19.24 %) [46]. In fact, the greater oxidative stability of the oils from Coratina cultivar is based on their high MUFA/PUFA ratio as well as the high content of polyphenols and other minor components, such as pigments [50].

**Table 1.**
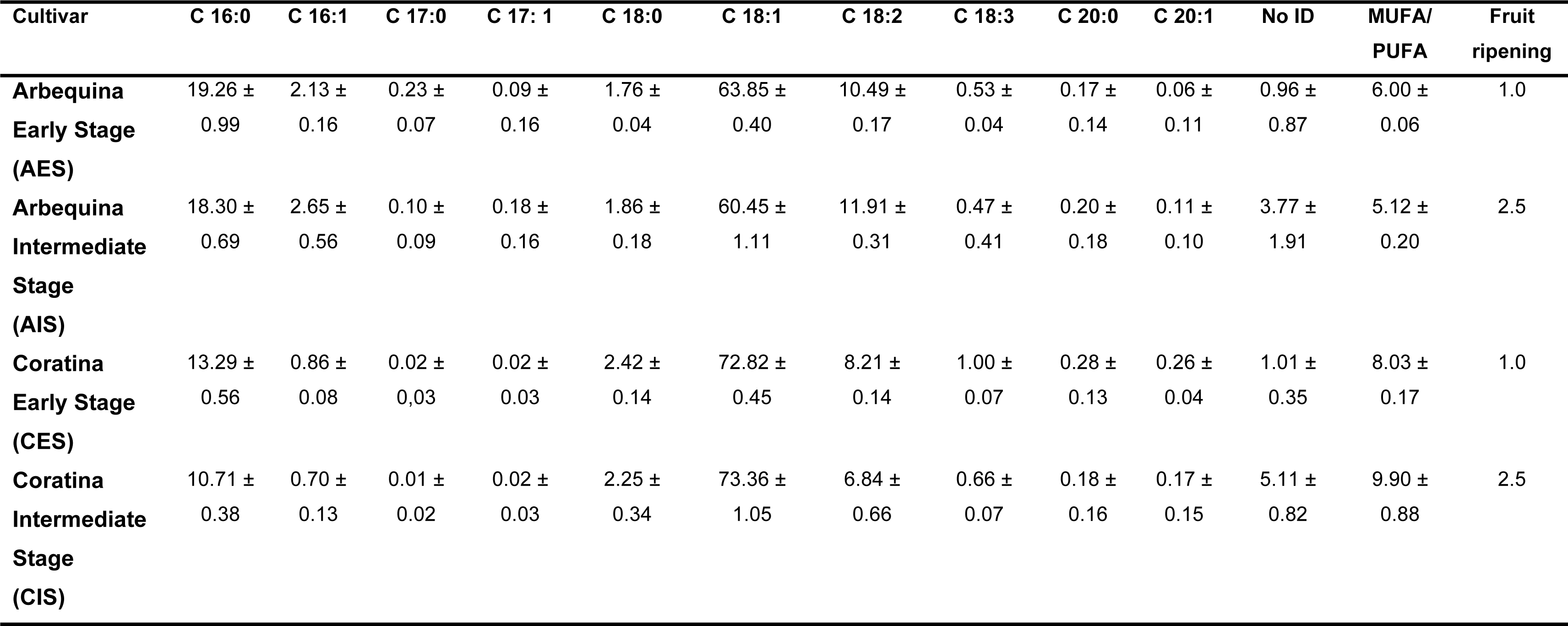
Profile of fatty acids (%) on Arbequina and Coratina and fruit ripening.

### Nitro-fatty acids formation depends on the cultivar and fruit ripening

Since endogenous levels of NO_2_-FA are very low compared to the formation of NO_2_-FA under acidic gastric nitration conditions, the present study analyzes the content of NO_2_-FA when olive oil was subjected to gastric acidic milieu. Under these conditions, the presence of NO_2_-OA, NO_2_-LA and NO_2_-cLA was detected being NO_2_-OA and NO_2_-cLA the most prevalent (Fig 1). While OA is the most abundant fatty acid in olive oil compared to cLA, it is less susceptible to nitration than NO_2_-cLA, which is the preferential unsaturated fatty acid substrate for nitration reactions during oxidative inflammatory conditions and digestion [51].

**Figure 1.**
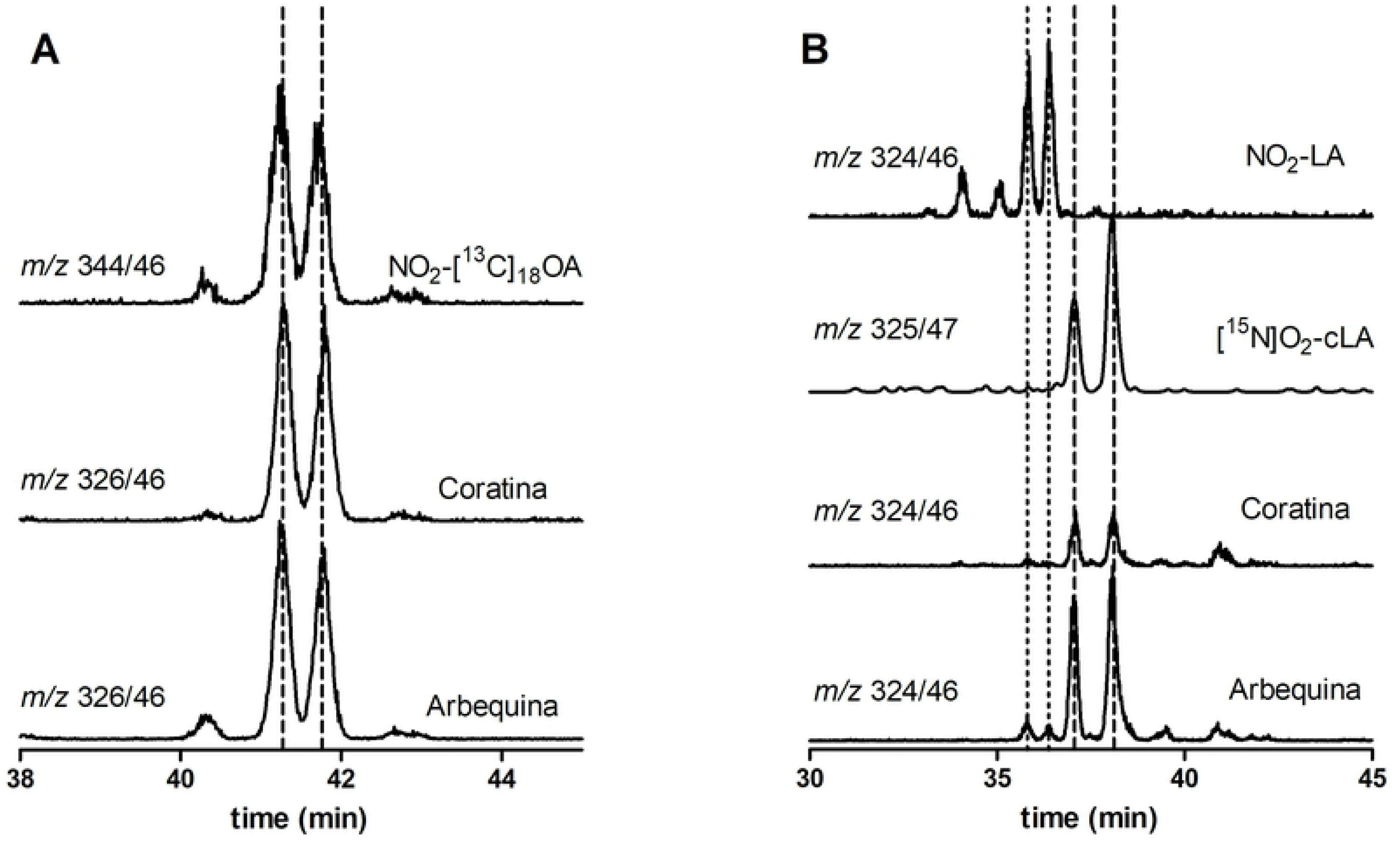
NO_2_-FA generation from VOO by *in vitro* digestion modeling. (A) The formation of NO_2_-OA in Arbequina and Coratina oils was determined following the MRM transition *m/z* 326/46 compared to the internal standard NO_2_-[^13^C_18_]OA (*m/z* 344/46). (B) The presence of NO_2_-LA and NO_2_-cLA in VOO gastric fluid was determined following the MRM transition *m/z* 324/46 compared to the internal standard [^15^N]O_2_-cLA (*m/z* 325/47) and external standard NO_2_-LA (*m/z* 324/46). Data shown is representative of at least 3 independent experiments.

Next, we compared the formation of the main detected NO_2_-FA: NO_2_-OA (Fig 1A) and NO_2_-cLA (Fig 1B) in Arbequina and Coratina during ES and IS conditions. As shown in Figs 2A and 2B, during the *in vitro* digestion of VOO the formation of NO_2_-OA and NO_2_-cLA in Arbequina was not dependent on the rip stage. However, the formation of NO_2_-FA in Coratina was related to rip stage, where maximal levels of NO_2_-cLA were observed at the IS (Fig 2B). The differences found on these two cultivars may be due to the effect of the MUFA/PUFA ratio, as show in Table 1. In addition, the presence of other minor components present in the olive oil which depend on a variety of factors, including cultivar and maturity index [39] may also contribute to the observed results. Olive oil mostly consist of triacylglycerols (TGAs, 98-99%), but there is a plethora of micro constituents present in VOO including phenolic compounds, that are relatively abundant and play an important role due to their antioxidant properties for food stability [52,53].

**Figure 2.**
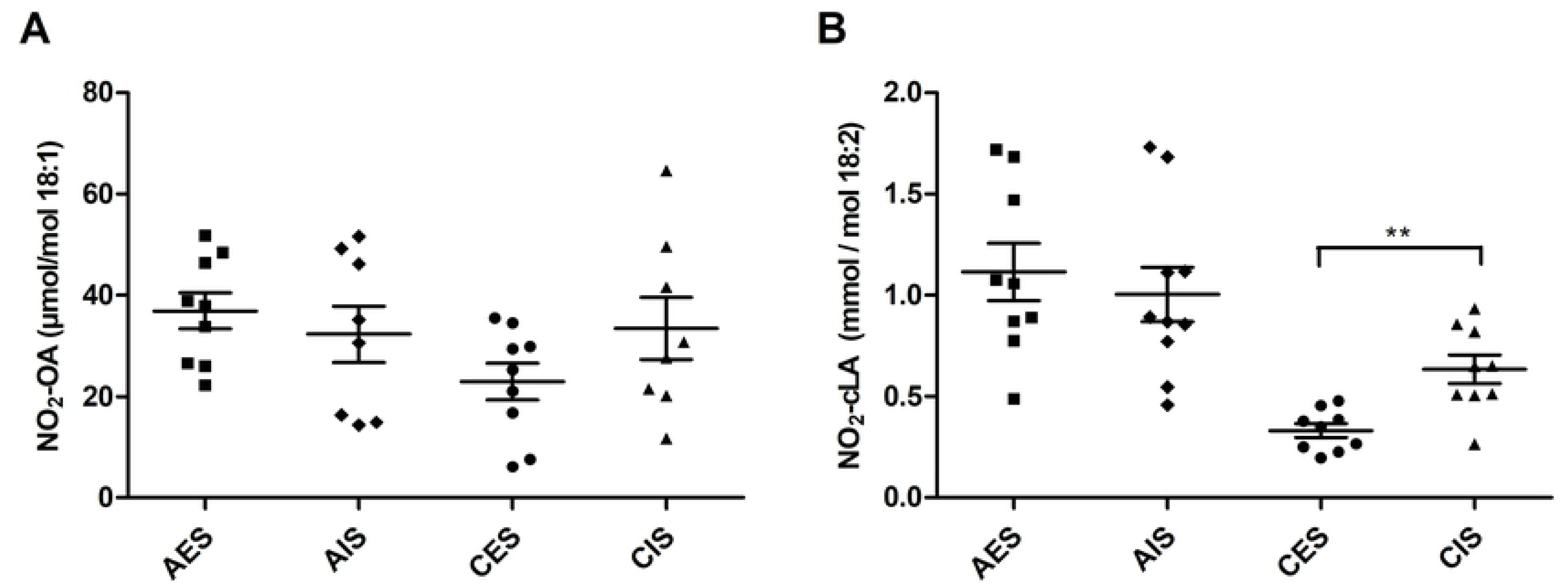
Effect of cultivar and rip stage in nitro-fatty acid formation. The formation under acidic gastric nitration conditions of (A) NO_2_-OA and (B) NO_2_-cLA from VOO in Arbequina, and Coratina during ES and IS conditions. Values are mean of three independent experiments. Differences were significant when p ≤ 0.05 (*) and p ≤ 0.01 (**).

### Polyphenols and pigments affect fatty acid nitration

Table 2 shows significant differences in the content of polyphenols between cultivars being the polyphenols content greater in Coratina than Arbequina. Within Arbequina cultivar, the ES exhibited greater total polyphenol content compared to the IS. In contrast, Coratina in IS increased these compounds 50% compared to ES. In the same line, other authors observed that the content of polyphenols in Arbequina was lower than Coratina [50] which is in line with the differences we observed on the MUFA/PUFA ratio. Five categories have been proposed to classify olive oils by their content of polyphenols: very high (> 600 g/Kg), high (450–600 g/Kg), medium (300–450 g/Kg), low (150–300 g/Kg), and very low (< 150 g/Kg) [54]. According to their content of polyphenols, Arbequina oils were mainly classified into category “very low” and Coratina oils into category “low”.

**Table 2.** Polyphenols, chlorophylls and carotenoids content of VOO.

| Compounds | Arbequina<br>Early Stage<br>(AES) | Arbequina<br>Intermediate<br>Stage<br>(AIS) | Coratina<br>Early Stage<br>(CES) | Coratina<br>Intermediate<br>Stage<br>(CIS) |
| --- | --- | --- | --- | --- |
| **Polyphenols**<br>(mg/Kg ) | 55.60 ± 5.15 | 34.17 ± 2.11 | 108.40 ± 10.02 | 216.27 ± 5.69 |
| **Chlorophylls**<br>(mg/Kg) | 4.49 ± 0.43 | 2.55 ± 0.47 | 29.74 ± 4.40 | 15.65 ± 3.22 |
| **Carotenoids**<br>(mg/Kg) | 3.73 ± 0.42 | 2.28 ± 0.27 | 17.37 ± 1.92 | 8.26 ± 1.29 |

In order to elucidate if the phenolic content can affect NO_2_-FA formation, polyphenols were extracted from different oils and fatty acid nitration was evaluated. Figs. 3A and 3B show that, in the case of Arbequina oil, the extraction of polyphenols reduce the amount of NO_2_-OA and NO_2_-cLA by approximately 70% in both maturation states. During digestion, the acidic environment in the stomach favors •NO_2_ production from NO_2_^-^, promoting a nitrative environment. Of relevance, fatty acid nitration depends on the surrounding O_2_ levels; while •NO_2_ can initiate lipid oxidation processes in the presence of O_2_, at low O_2_ levels fatty acid nitration predominates over oxidation (*e.g.* lipoperoxidation) [25,55] (Scheme 1).

**Figure 3:**
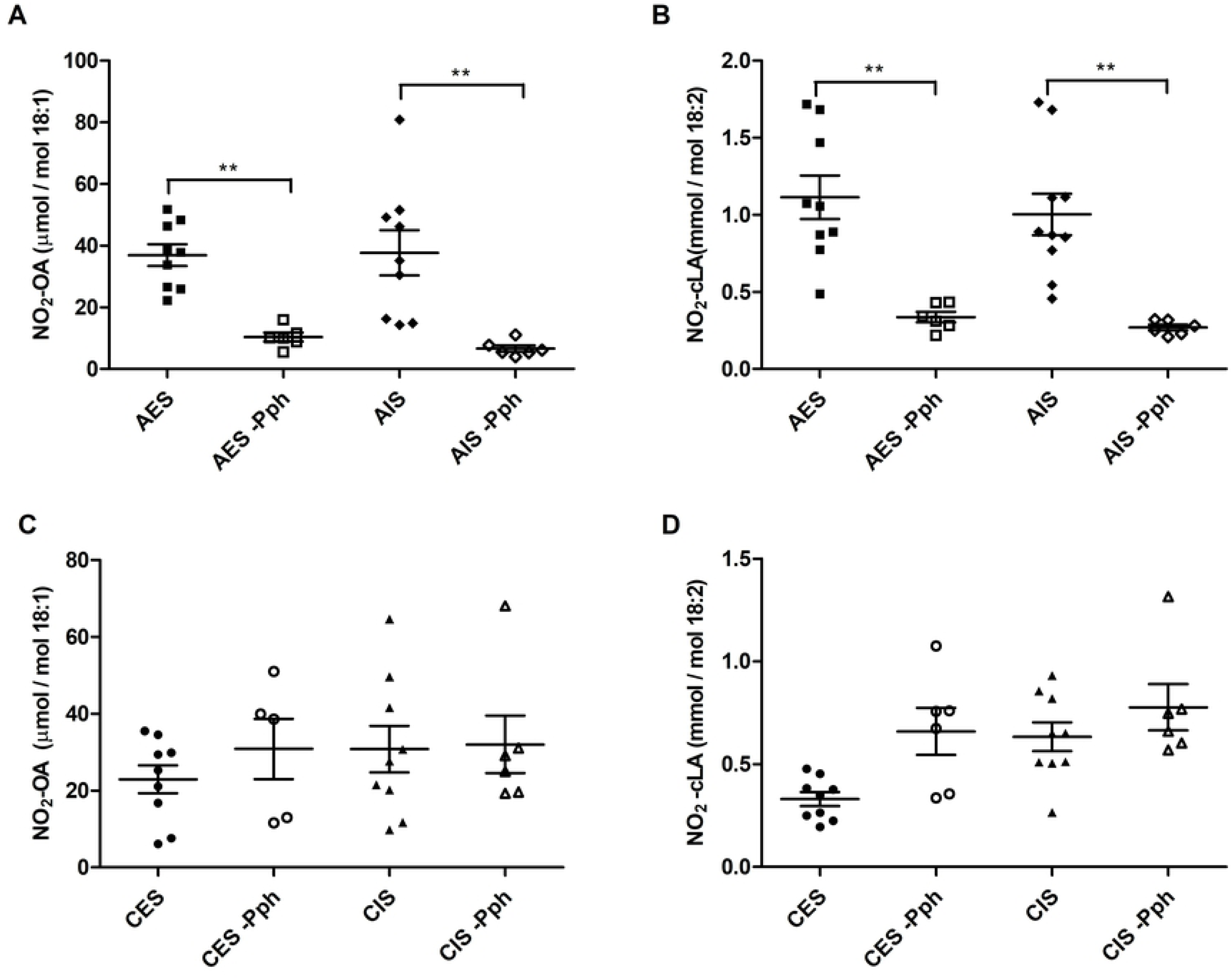
Effect of polyphenols in NO_2_-FA formation in VOO. (A) NO_2_-OA and (B) NO_2_-cLA in Arbequina; (C) NO_2_-OA and (D) NO_2_-cLA in Coratina. Values are means of three independent values. Differences were significant when p ≤ 0.05 (*) and p ≤ 0.01 (**).

Based on the results reported here, we propose that polyphenols play a key role during digestion protecting fatty acids from oxidation and, in turn, promoting nitration pathways to form products such as NO_2_-FA. On the other hand, in the absence of polyphenols the oxidation pathways would be favored, where •NO_2_ initiate and propagate fatty acid oxidation, thus decreasing fatty acid nitration. In Coratina (Figs 3C and 3D), where the content of polyphenolic compounds is higher than in Arbequina, the absence of these molecules did not affect the formation of NO_2_-FA. There are other bioactive constituents in olive oils such as tocopherols and pigments, that can explain the observed differences between Coratina and Arbequina [56]. Pigments, constituted by CAR and CHL derivatives determine the color and are related to VOO quality due to their relationship with freshness, nutritional and antioxidant properties [15,16]. Many factors, such as cultivar, geographic origin, maturity of olives, climate and storage conditions influence pigments content [56]. Previous studies shown that in Arbequina oils the content of polyphenols, CAR and CHL is lower than Coratina reducing the oxidative stability of theses oils [50]. In line with these results, we analyzed CHLs and CARs content in these two cultivars. As shown in table 2, Coratina oil contains the highest levels of these pigments when compared with Arbequina. These results linked to the low MUFA/PUFA ratio, explain that Arbequina oils are more susceptible to oxidation, decreasing nitration of fatty acids especially in the absence of polyphenols. On the other hand, Coratina oils have the higher oxidative stability that is based on their better fatty acids profile in addition to their polyphenols and pigments content. Moreover, the absence of polyphenols does not affect to formation of NO_2_-FA, due to the highest content of pigments compared to Arbequina, overall protecting oil from oxidation by favoring the fatty acid nitration pathway (Scheme 1)

We conclude that a) VOO serves as precursors of NO_2_-FA and b) bioactive constituents in olive oils such as polyphenols and pigments could protect VOO from lipid oxidation favoring fatty acid nitration (Scheme 1). Based in our results, we postulate that the composition of VOO assures the formation of NO_2_-FA and their health beneficial properties.

## Acknowledgment

This work was supported by grants from ANII-Innovagro FSA 1 2013 1 12622 and CSIC-Grupos N° 536 to HR, and CSIC I+D to AT. BSC was supported by a postdoctoral fellowship by ANII-Uruguay. We are grateful to Juan José Villamil and David Bianchi for his support in processing samples and olive oil elaboration and to Juliana Bruzzone and Cecilia Martínez for her contribution in the quality olive oil analysis

